# Dynamic connectivity maps of pericytes and endothelial cells mediate neurovascular coupling in health and disease

**DOI:** 10.1101/830398

**Authors:** Tamas Kovacs-Oller, Elena Ivanova, Paola Bianchimano, Botir T. Sagdullaev

## Abstract

Functional hyperemia, or matching blood flow to activity, is spatially accurate to direct the oxygen and nutrients to regionally firing neurons. The underlying signaling mechanisms of neurovascular coupling remain unclear, but are critical for brain function and establish the diagnostic power of BOLD-fMRI. Here, we described a mosaic of pericytes, the vasomotor capillary cells in the living retina. We then tested if this symmetric net of pericytes and surrounding neuroglia predicted a connectivity map in response to sensory stimuli. Surprisingly, we found that these connections were not only discriminatory across cell types, but also highly asymmetric spatially. First, pericytes connected predominantly to other neighboring pericytes and endothelial cells, and less to arteriolar smooth muscle cells, and not to surrounding neurons or glia. Second, focal, but not global stimulation evoked a directional vasomotor response by strengthening connections along the feeding vascular branch. This activity required local NO signaling and occurred by means of direct coupling via gap-junctions. By contrast, bath application of NO or diabetes, a common microvascular pathology, not only weakened the vascular signaling but also abolished its directionality. We conclude that the discriminatory nature of neurovascular interactions may thus establish spatial accuracy of blood delivery with the precision of the neuronal receptive field size, and is disrupted early in microvascular disease.

**Highlights:**

- Within a structurally symmetric mosaic, pericytes form discriminatory connections
- Pericyte connectome tunes with a precision matching a neuronal receptive field
- Focal but not global input evokes a vasomotor response by strengthening the gap-junction mediated signaling towards a feeding vascular branch
- Disrupted functional connectivity map triggers loss of the functional hyperemia in diabetic neuropathy

## Introduction

Local changes in neural activity evoke a vascular response that is spatially restricted to the activated region. The ubiquitous nature of the microvasculature and the diversity of routes through which blood can be distributed make this task challenging. Furthermore, blood perfusion must be shifted or expanded to accommodate for changing activity patterns. To accomplish changes in demand, functional hyperemia must involve at least three events with a spatial precision that enables discrimination of the active site from its resting neighbor: 1) sensing local changes, 2) transmission of vasoactive signals along the supplying vascular branch, culminating in 3) vasomotor response that directs blood to the active region. The mechanisms responsible for this spatial accuracy are not clear, but are critical for brain function and establish the diagnostic power and precision of BOLD-fMRI^1^.

The strategic location of capillaries within synaptic layers where neurotransmitters are released, supports their function as both sensors and responders to neural activity^2, 3^. Here, pericytes, the only contractile cells along a vast capillary network may fulfill both of these roles. Contrary to long held beliefs, pericytes express smooth muscle actin^4^, which enables capillary diameter changes in response to electrical, pharmacological and sensory stimuli^2, 5–7^. In light of these observations, it is surprising that the vasomotor response due to sensory stimulation is not observed initially at proximal capillary regions, but rather in larger vascular branches, away from the sites of neural activity^6, 8^. While further verification is needed, this intriguing spatial segregation is consistent with a primary role of larger precapillary regions in blood supply^9, 10^. More importantly, on a temporal scale, it suggests the existence of a presently unknown signaling mechanism that enables rapid propagation of a vasoactive signal from the capillary region towards the feeding vascular branch^11^. Direct gap junction-mediated communication among pericytes and other vascular elements play a key role in vasomotor response propagation^6, 12, 13^. Indeed, in the isolated retina vasculature, electrotonic pulses propagate radially along the network of pericytes and endothelial cells^14^. However, this radial spread is inconsistent with the directional nature of the vasomotor response occurring in the intact brain. Following focal stimulation, the vasomotor response tends to propagate stronger in the direction of the feeding precapillary region, a phenomenon widely observed in multiple systems, including retina^9^, olfactory bulb^15^, cerebral cortex^16^, and skeletal muscles^17^. These studies expose the following key questions: a) what is the nature of cell-to-cell interactions between the different regions of the vascular tree that mediate the spatio-temporal precision of a vasomotor response, b) how changes in local neural activity induce directional bias within a radially-coupled vascular synthicium, and c) how vascular signaling accommodates for changes in sensory modalities?

Recent characterization of the “vascular relay” in the retina, a specialized vascular region along a capillary-arteriolar branch with distinct distribution of Cx43-containing gap junctions across endothelial cells and pericytes, established structural foundation for vascular cell connectivity^13^. Here, in retina wholemount, a self-contained brain structure with a defined vasculature, we experimentally tested the hypothesis that the spatial accuracy of vasomotor response is driven by a precise and discriminatory connectivity map among vascular cells - pericytes and endothelial cells. We further probed whether, and ultimately how, these connectivity maps dynamically shifted to accommodate to the changing sensory input. To accomplish this, we used a combinatorial approach of light simulation, direct cell-to-cell tracer coupling, multiphoton imaging of calcium activity, and the measurements of vasomotor response and blood flow. Use of intact retinal wholemount preparation has allowed us to rigorously combine natural light stimulus with selective pharmacological tools to study fundamental properties of vascular signal transmission with high temporal and spatial resolution. We then used this model system to evaluate how its impairment may contribute to diabetic retinopathy, a common vascular pathology in diabetic patients^18, 19^.

## Results

### Pericytes form a mosaic in the retina microvasculature

Functional hyperemia is thought to rely on coordinated activity across a broad vascular network, directing blood to active regions. Since a vasomotor response will depend on placement of the vasoactive elements, the knowledge of pericyte distribution across a broad capillary network is needed. In NG2-DsRed mice, all vessels, including arterioles, veins and capillaries, were readily identified by fluorescently-labeled mural cells. In these mice, we combined a set of anatomical features with a selective uptake of NeuroTrace 500/525 (NT500) assay to unambiguously distinguish pericytes from smooth muscle cells (Figure 1A-C)^19, 20^. In each vascular layer within a 300×300 μm retinal patch, pericyte cell bodies were notated (Figure 1D). For each vascular lamina, two sets of measurements were calculated: the shortest route along the vessel (SRAV), and the nearest-neighbor distance (NND) between pericytes, regardless of their capillary residence (Figure 1E). Across all vascular layers, we found that pericytes were distributed regularly (Gaussian goodness of fit R2 = 0.93 - 0.99, n = 16 mice). This regularity was maintained even in the superficial vascular layer (R2 = 0.93), despite areas occupied by large precapillary regions (Figure 1F). Overall, inter-pericyte distances decreased from superficial to deep vascular layer, consistent with increased capillary density. As an exception, SRAV distance was longest in the intermediate layer, likely due to inherent tortuosity of the capillary network. For pericyte distribution, conformity ratio (CR), a robust measure for structural mosaics (see Methods), was significantly above either Ready-Reckoner thresholds (2.3 for n = 50 samples)^21^ or randomly generated values, drawn from the same data set (SRAV: 3.63 ± 0.76 vs. 1.74 ± 0.27, P < 0.001; NND: 4.47 ± 0.72 vs. 1.71 ± 0.19, P < 0.001, paired T-TEST, n = 48 ROIs in n = 16 mice for each, Figure 1G). Strikingly, NND conformity ratios were higher compared to SRAV (superficial: 4.31 ± 0.72 vs. 3.58 ± 1.13, P = 0.019, intermediate: 4.75 ± 0.77 vs. 3.64 ± 0.42, P < 0.001; deep: 4.36 ± 0.64 vs. 3.67 ± 0.59, P < 0.001 paired T-TEST n = 16 mice for each). Thus, the pericyte mosaic was not simply driven by direct pericyte-to-pericyte connectivity along the same vascular branch, suggesting that their placement is optimized for volume regularity across the retina.

**Figure 1.**
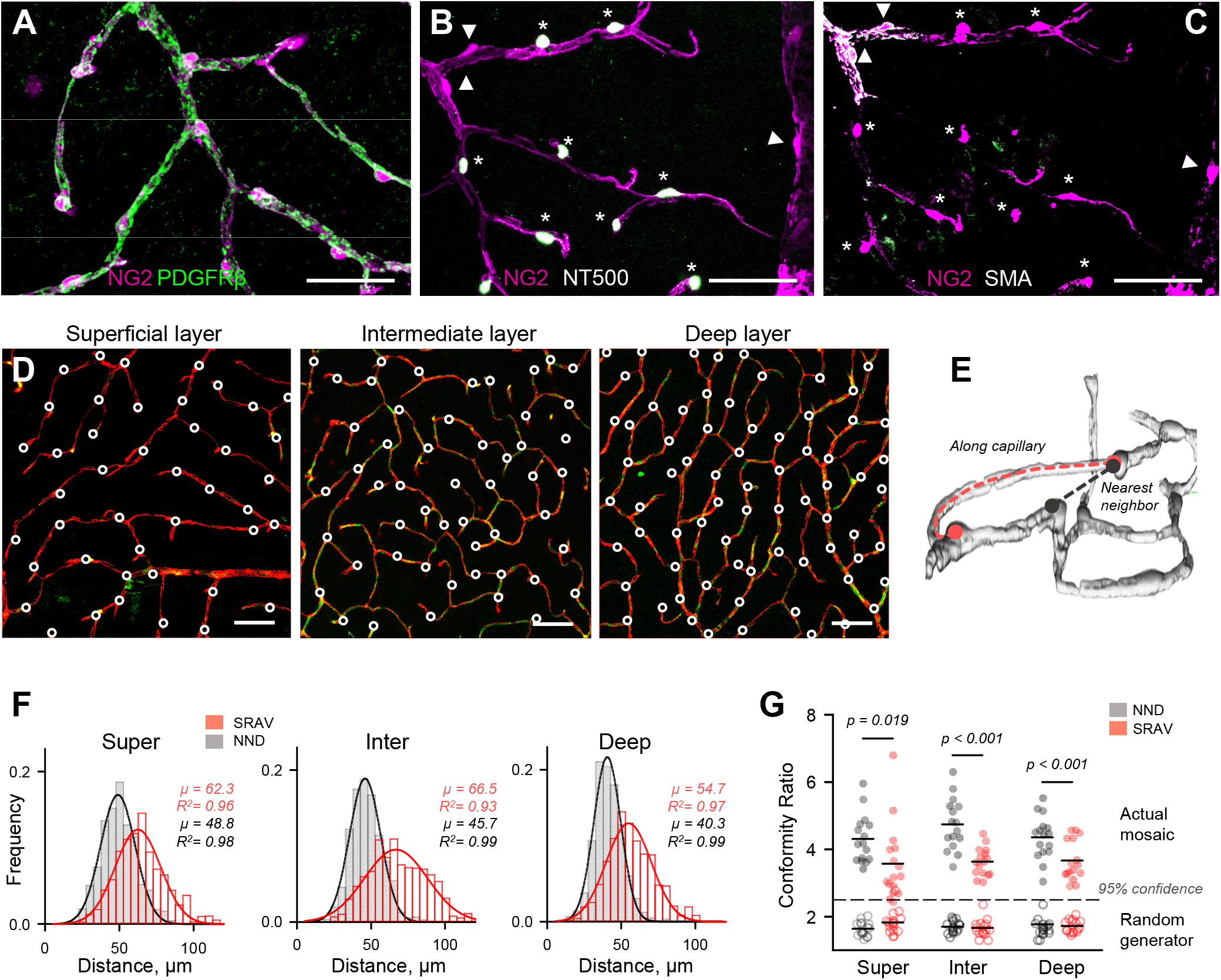
Mosaic or pericytes in the retinal vasculature. **(A-C)** Pericytes (asterisks) were distinguished from other mural cells (arrowhead) using a selective uptake of NT500. **(D)** Distribution of pericytes across three retinal vascular layers. For visualization and a follow up analysis, each circle was placed over the pericyte cell body. Scale 50 μm. **(E)** Illustration showing how both the inter-pericyte shortest route along the vessel (SRAV) and the nearest neighbor distance (NND) were measured. **(F)** Distance frequency histograms in each vascular layer. Mean distances (*μ*) and the Gaussian goodness of fit (*R*^2^) for each distribution are indicated to the right. **(G)** Conformity Ratios (mean/SD) for NND and SRAV. Dotted line represents a 95% confidence threshold over randomly-generated values drawn from the same distribution. Each point in the plot represents a mean value within individual animal (n = 16 mice) with a combined number of measures n = 3647. *P*-values are from one-way ANOVA with Tukey’s post-hoc test.

### Vascular cells establish discriminatory GJ connections with each other but not with neurons or glia

What are the mechanisms of vasomotor response propagation along the vascular tree? The answer to this question will depend not only on spatial distribution of vasoactive elements, but also on the nature of underlying cell-to-cell communication. Direct and rapid communication among neurovascular elements via gap junctions (GJ) is a key event in vasomotor response propagation; GJ block completely abolished response propagation, but not its initiation^6^. To establish a neurovascular connectivity map, or “neurovascular connectome” of the retina, we used single-cell injection of gap junction-permeable probe Neurobiotin (NB) and traced all cells that were coupled to the injected cell^22^.

In the retina wholemount, we targeted individual mural, endothelial cells, astroglia and neurons. To identify each cell we relied on a set of characteristic anatomical features, molecular and immunohistochemical markers (Figure 2D). As illustrated in Figure 2C,E, 15 minutes following NB injection, a “chain” of GJ-coupled cells could be revealed (Suppl.Vid.1). To unambiguously identify the injected cell, we supplemented the NB-containing intracellular solution with a larger, GJ-impermeable Alexa dye (Figure 2B-D). Furthermore, the NB spread was precluded by meclofenamate (MFA, 40 μM), a selective GJ blocker (Figure 2E,G). In MFA-treated retina, both Alexa and NB were restricted to the targeted cell (Extended Data S2). Together, this evidence confirms that NB was selectively infused into a targeted cell and its spread occurred through GJs, and not by nonspecific NB uptake.

**Figure 2.**
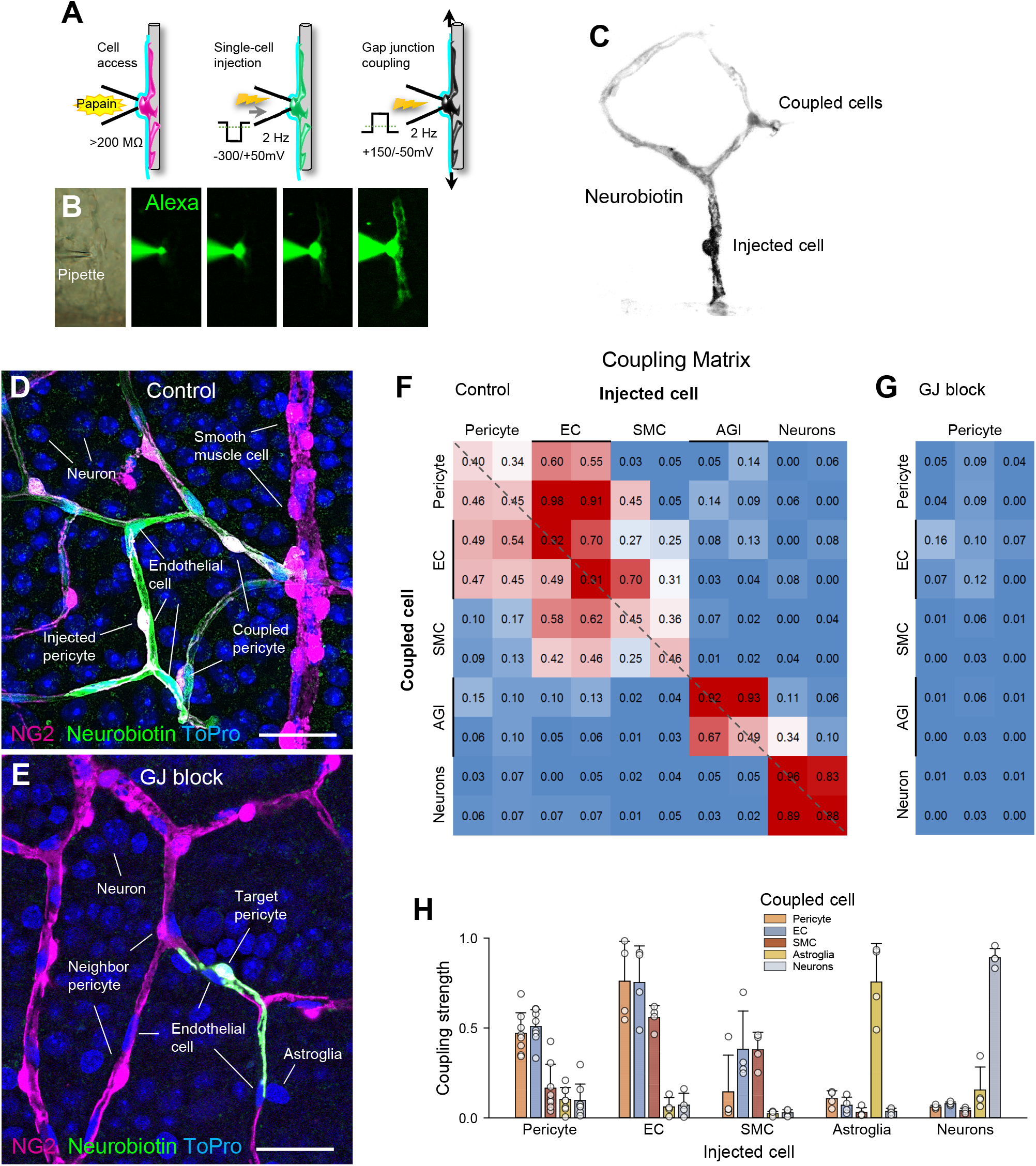
Coupling connectome of the neuro-glia-vascular unit. **(A)** Illustration of the experimental design to target and trace GJ-mediated connections of an identified cell using a combination of probes with different size/permeability. **(B)** With a patch-clamp pipette, large Alex488 dye and small Neurobiotin tracer are co-injected into the target cell. Using this assay, distinct cells can be selectively targeted and their connections traced (Suppl. Fig.1). **(C)** Appearance of Neurobiotin spread from the targeted cell is revealed with an anti-streptavidin antibody. **(D)** Confocal reconstruction of the tracer spread across GJ-coupled cells in the retinal wholemount (Suppl. Vid.1). **(E)** Treatment with a GJ blocker (meclofenamate, 40 μM) precludes tracer spread. Scale 25 μm. **(F, G)** Cell-type specific connectivity matrices. In this heatmap matrix, the injected cell type is shown on top, with its coupled neighbor below. For each experimental condition, samples from 4-6 individual mice are shown. The number in each box represents coupling strength between the target cell and its four neighbors. Higher value and a warmer color indicates stronger coupling. **(H)** Summary of connectivity strength for individual pairs across 6-8 mice per each condition..

Next, for each cell type, coupling strength (CS), a measure of NB spread (Methods) was calculated to yield a coupling matrix of the retinal neurovascular unit (Figure 2F,G). The analysis of coupling matrices revealed a number of insights. The majority of connections were homotypic and highly discriminatory across cell types; vascular cells coupled to other vascular cells, astroglia to astroglia, and neurons to neurons. Here, due to lack of heterotypic GJ-coupling, we simplified the approach and treated all neuronal cells as a single group, without further discrimination. Pericytes were coupled predominantly to other neighboring pericytes (CS = 0.47 ± 0.11) and endothelial cells (0.51 ± 0.09, P = 0.24) and significantly less to arteriolar smooth muscle cells (0.16 ± 0.13, P < 0.001), glia and neurons (0.10 ± 0.06 and 0.10 ± 0.09, respectively, P = 0.001, multi-comparisons ANOVA, Tukey post-hoc, n = 8 mice for each pair). Interestingly, smooth muscle cells on the arterioles were among the least coupled cells (Figure 2 F,H and Suppl.Figure 1), evidence for a mechanism, limiting the changes in blood supply beyond the activated region and thus improving spatial accuracy of functional hyperemia.

### Pericyte and endothelial cell connectivity maps shift in response to changing sensory input

Having established a spatial mosaic of underlying vasomotor elements, pericytes and their exclusive connectivity maps, we next used the living retina preparation and its natural stimulus light, to test how these interactions change with sensory input. In the brain, neurons are highly tuned to preferential modalities. In particular, retinal neurons tend to respond strongly to stimuli limited to their receptive field size. We hypothesized that this activity would also translate to vascular map activation. If true, this would establish a theoretical upper resolution limit for BOLD-fMRI. To accomplish this, we compared vascular cell coupling under two experimental conditions: a) full-field flickering light (4Hz) and b) spot flickering light (4Hz, 150 μm diameter, that approximates the excitatory center of a retinal ganglion cell receptive field)^23, 24^. To ensure that equal amounts of light were delivered under both approaches, a stationary background was maintained at a mean value of full-field stimulus around a flickering spot center. For easier access and parfocal view of diverse vascular branches, pericytes in the superficial layer were targeted. Nevertheless, NB spread was evaluated across all vascular layers (Extended Data S2). For consistency across multiple samples, we injected pericytes 3-4 branch points away from the feeding artery (~300 μm, Figure 3A-D), thus allowing sufficient space in both up and downstream directions of the vascular tree. We then measured the coupling strength and the directionality index (DI), the measure of upstream vs. downstream bias (Figure 3F, Methods). DI values >1 indicate bias towards the feeding artery. We found that under full-field light stimulus, pericytes were coupled to other pericytes and endothelial cells with CS = 0.47 ± 0.12 and 0.51±0.09 (n = 8 mice each, Figure 3E), respectively. No directional bias was evident (DI = 0.77 ± 0.21), consistent with an even distribution of the vasomotor elements. In stark contrast, spot light stimulus significantly increased coupling strength over full-field stimulation (CS = 0.79 ± 0.11, P < 0.001, ANOVA with Tukey’s post-hoc, n = 8 mice, Figure 3B). Surprisingly, this was not driven by a symmetric increase in coupling in both directions from the injected pericyte, as would be predicted from their structural regularity (Figure 3H-G). Instead, focal sensory input strengthened connections along the shortest path towards a vascular branch feeding the active site with DI = 2.0 ± 0.4 (P < 0.001, ANOVA with Tukey’s post-hoc, n = 8 mice), establishing a characteristic “footprint” of a directional vasomotor response. Again, both coupling and directionality were blocked by MFA (40 uM, CS = 0.07 ± 0.06; DI = 0.84 ± 0.15, n = 6 mice).

**Figure 3.**
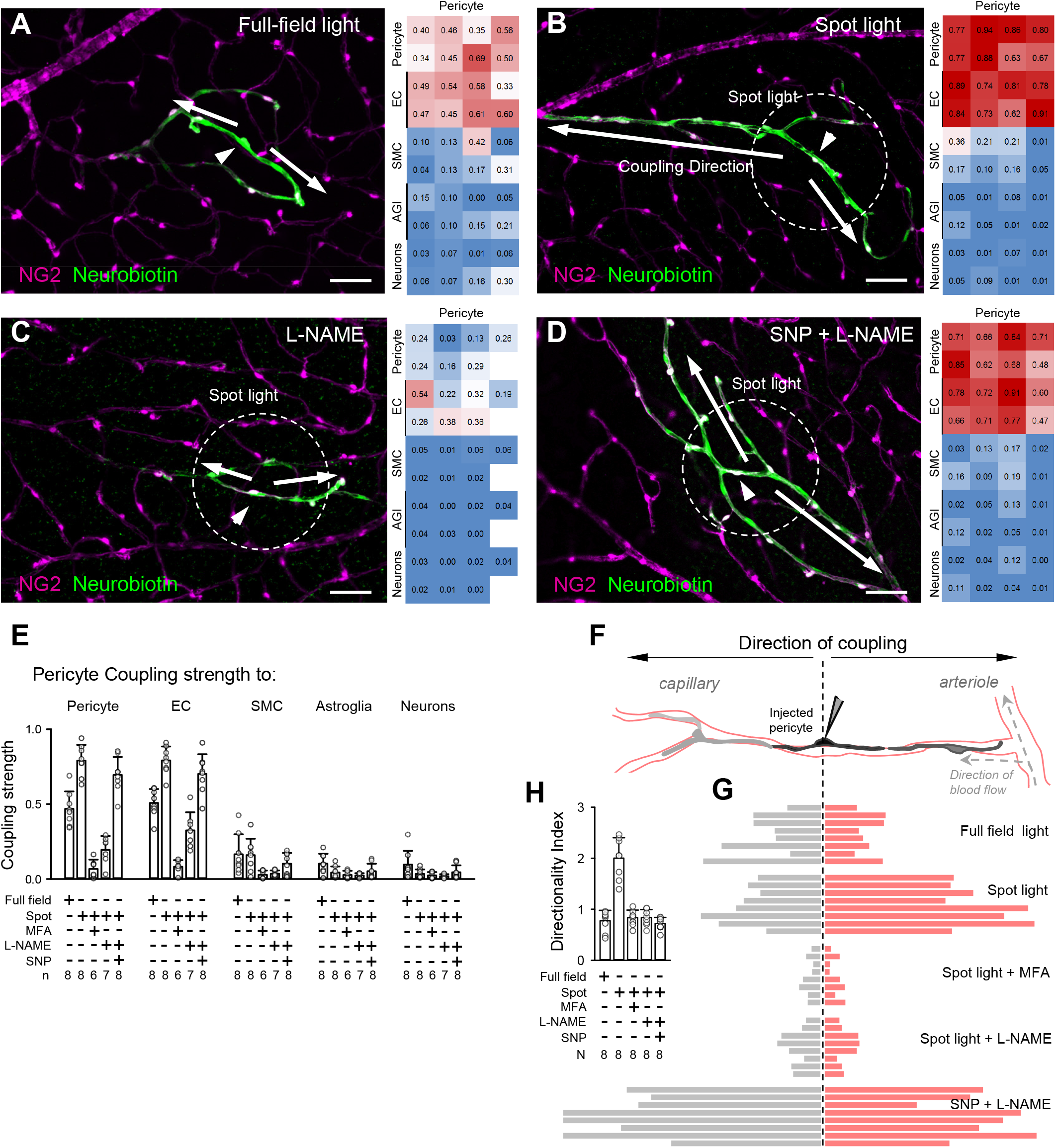
Strength and directionality of vascular coupling is driven by focal sensory input and is mediated by NO-signaling. **(A)** Spatially broad sensory stimulation (full-field light) produces weak pericyte-to-pericyte and pericyte-to-endothelial cell coupling. **(B)** Focal sensory stimulation under capillary area induces strong vascular coupling along the capillary branch, biased towards the feeding arteriole. **(C)** Blockade of NO-signaling abolished increased coupling strength induced by a focal sensory stimulation (150 um light spot). **(D)** exogenous application of NO-donor significantly increases coupling strength, but fails to restore its directionality. Scale bars 50 μm. **(E)** Summary of pericyte cell coupling strength under each experimental condition. **(F)** Experimental design and assessment of vascular cell coupling directionality. **(G)** Vascular cell coupling strength up and downstream from the injected pericyte (dashed vertical line). Each bar pair (left and right) is from individual mice under each experimental condition (n = 8 mice per each group). **(H)** Summary of pericyte cell coupling directionality under experimental conditions._28_

### Directionality of vasomotor response is driven by local NO signaling

Our findings provide a direct evidence for a dynamic connectivity map utilizing a structurally rigid vascular framework^25^, suggesting a potential role in spatial accuracy of functional hyperemia. To further test this, we determined whether the increase in vascular cell coupling was indeed driven by neural activity. Synthesis of NO is a key event in the neurovascular coupling^5, 7^. We therefore hypothesized that blocking nitric oxide synthase (NOS), an enzyme responsible for activity-induced NO production, would diminish vascular coupling. Consistently, during spot light stimulation, application of L-NAME (100 μM), a broad-spectrum NOS inhibitor, significantly reduced vascular coupling compared to spot light alone (0.20 ± 0.09 vs. 0.79 ± 0.11, P < 0.001, ANOVA with Tukey’s post-hoc, n = 7 and 8 mice, respectively). It also abolished directionality of vascular cell coupling (Figure 3 F-H, 0.84 ± 0.16 vs. 2.0 ± 0.4, P < 0.001, ANOVA with Tukey’s post-hoc, n = 7 and 8 mice, respectively). In the presence of L-NAME, vascular cell coupling was also significantly lower than under full-field sensory stimulation (0.2 ± 0.09 vs. 0.47 ± 0.12, P < 0.001, ANOVA with Tukey’s post-hoc, n = 7 and 8 mice, respectively), suggesting sustained coupling under suboptimal sensory stimulation. However, this activity was not sufficient to evoke directional response. Was this strengthened directionality of vascular coupling driven by simply an overall increase in NO production due to optimized neuronal stimulation, or did it rather require spatially defined *local* increase in NO? To address this, we next measured the magnitude of NB spread in response to spot light stimulation in the presence of both L-NAME, to block local NO production, and bath applied sodium nitroprusside (SNP, 100 μM). Bath application of NO significantly increased the coupling strength (Figure 3G, CS = 0.7 ± 0.12 vs. 0.2 ± 0.09 in L-NAME alone, P < 0.001, ANOVA with Tukey’s post-hoc, n = 8 and 7 mice, respectively). However, this was not accompanied by the characteristic directional profile of the NB coupling in response to spatially restricted stimulus (spot DI = 2.0 ± 0.4 vs. bath SNP DI = 0.72 ± 0.12, P < 0.001, ANOVA with Tukey’s post-hoc, n = 8 mice each), suggesting that local NO production was necessary for directional coupling.

Next, we tested whether locally applied NO will be sufficient for directional vasomotor response propagation. Indeed, focal light stimulation produced a vasomotor dilatory response that propagated preferentially upstream feeding vascular branch, but not downstream or collateral capillary regions (Figure 4A,B). This finding is consistent with an earlier established GJ-mediated coupling map (Figure 3G). Block of neurovascular coupling with L-NAME resulted in a loss of directional vasomotor response to spot light stimulation. Under full-field illumination, we focally stimulated a small area around individual pericyte, by puffing SNP using a patch pipette. To reduce the spatial spread of SNP, the pipette was positioned downstream to the vascular branch relative to perfusion flow. Similar to a spot light flicker, local puff of SNP produced directional vasomotor activity (Figure 4; note dilation in ROIs 1 and 2, but not 3 or 4). This directional response was lost during consequent bath application of SNP, resulting in broad vascular dilation in all vascular branched (Figure 4B, lower panels). This may provide a mechanistic basis of an earlier *in vivo* observation, showing that in the presence of NOS inhibitors, global NO supply failed to recover functional hyperemia of the cerebellar blood flow^26^. Together, these data indicate that spatially accurate stimulation is responsible for directionality of both cellular connectivity and the vasomotor response propagation.

**Figure 4.**
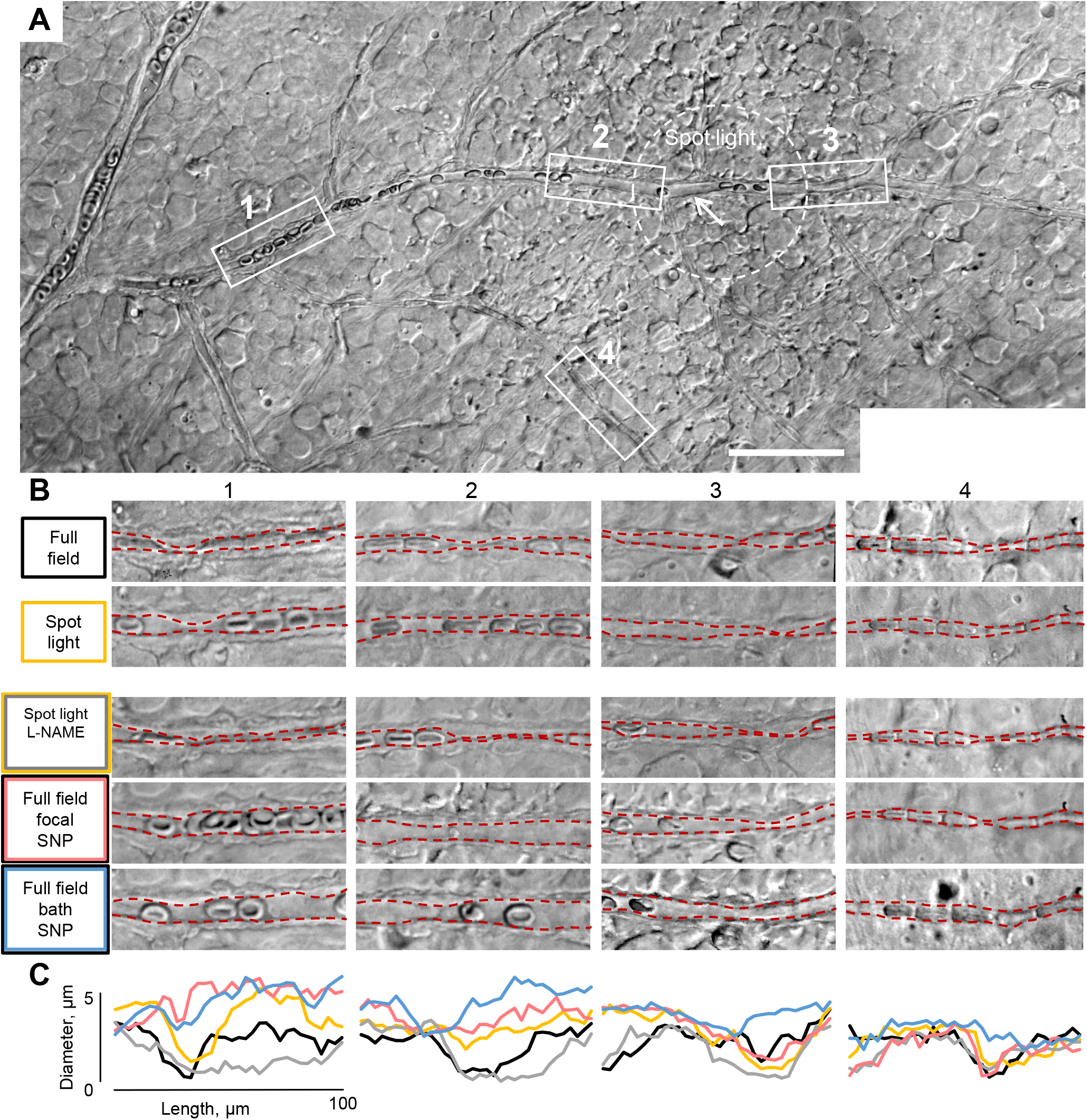
Directionality of light-induced vasodilation is driven by local, but not global NO signaling. **(A)** In the retina wholemount, vascular diameter changes were monitored at 100 μm long ROIs (boxed) below, above the site of stimulation and at the collateral branch (arrow). **(B)** Magnified ROIs under indicated experimental conditions. **(C)** Volumetric analysis of vascular diameter changes at each ROI. Largest vasodilation was observed in the direction toward the feeding branch (ROIs 2 and 1) in response to either flickering spot of light (150 μm, 4 Hz), or focal SNP puff (100 μM), but not during global light stimulation or in the presence of L-NAME (100 μM).

### Timing of Ca2+ change and vasomotor response is contractile cell type specific

Cellular basis of the vasomotor response dynamics remains controversial. This is in part due to a limited ability to control the activation site and timing, along with inaccurate vascular diameter measurements *in vivo* ^8, 9^. To overcome these limitations, we used freshly dissected retina wholemounts from NG2-Cre-GCamp6f mice. While in the earlier study, tamoxifen induction of Cre resulted in targeted expression of GCamp6f in mural cells exclusively^8^, in our mice, constitutive activity of the NG2 promoter resulted in GCamp6f expression in all retinal cells, including neurons, glia and vasculature. Next, we crossed our NG2-Cre-GCamp6f-EGFP mice with an NG2-DsRed line to label vasculature. Ubiquitous expression of GCamp6f (green fluorescence) and restricted expression of DsRed in the mural cells (magenta fluorescence) enabled us to simultaneously assess Ca^2+^ dynamics in the entire neurovascular unit and conduct accurate volumetric analysis of vasomotor response. To achieve precise timing and mimic local neuronal activation, individual pericytes were directly depolarized by a patch electrode. In NG2-Cre-GCamp6f-DsRed retina, we stimulated a capillary pericyte (2 ms, 10 μA) while imaging GCamp6f and vascular diameter under a two-photon microscope (Figure 5A, location 5). Short electric stimulation resulted in pericyte activation, followed by local transient activation of Muller cells (dashed circle marks the limit of the Muller cell activation) and propagation of Ca^2+^ wave through the vascular branch. DsRed-expressing contractile cells along the vascular branch were identified (labels in Figure 5A) and temporal calcium dynamics with vasoconstriction were measured at these locations at 15 frames per second (Figure 5B). The Ca^2+^ increase was almost instantaneous in the vascular branch leading towards the feeding artery (locations 1-4), but did not propagate much towards the vein (location 6). In spite of simultaneous Ca^2+^ increase, vasoconstriction, initiated by Ca^2+^, was significantly faster and stronger in smooth muscle cells (location 1-2) in comparison to pericytes (location 3-4). Again, propagation of both Ca^2+^ wave and vasomotor response relied on vascular GJs and was abolished by 40 uM meclofenamate (shown by the red line in Figure 4B, locations 1 and 3) without affecting Ca^2+^ rise and vasoconstriction at the targeted pericyte (location 5). To quantify temporal kinetics of the Ca^2+^ wave and vasomotor response as well as their relation to each other, we used the following parameters: peak amplitudes of Ca^2+^ increase and vasomotor response (ΔFmax/F and ΔDmax/D), time from stimulation to 10% and 90% of Ca^2+^ and vasoconstriction response. Figure 4C shows corresponding time frames for both smooth muscles and a pericyte. Both Ca^2+^ increase and vasomotor response were delayed in the pericyte in comparison with the smooth muscle cells.

**Figure 5.**
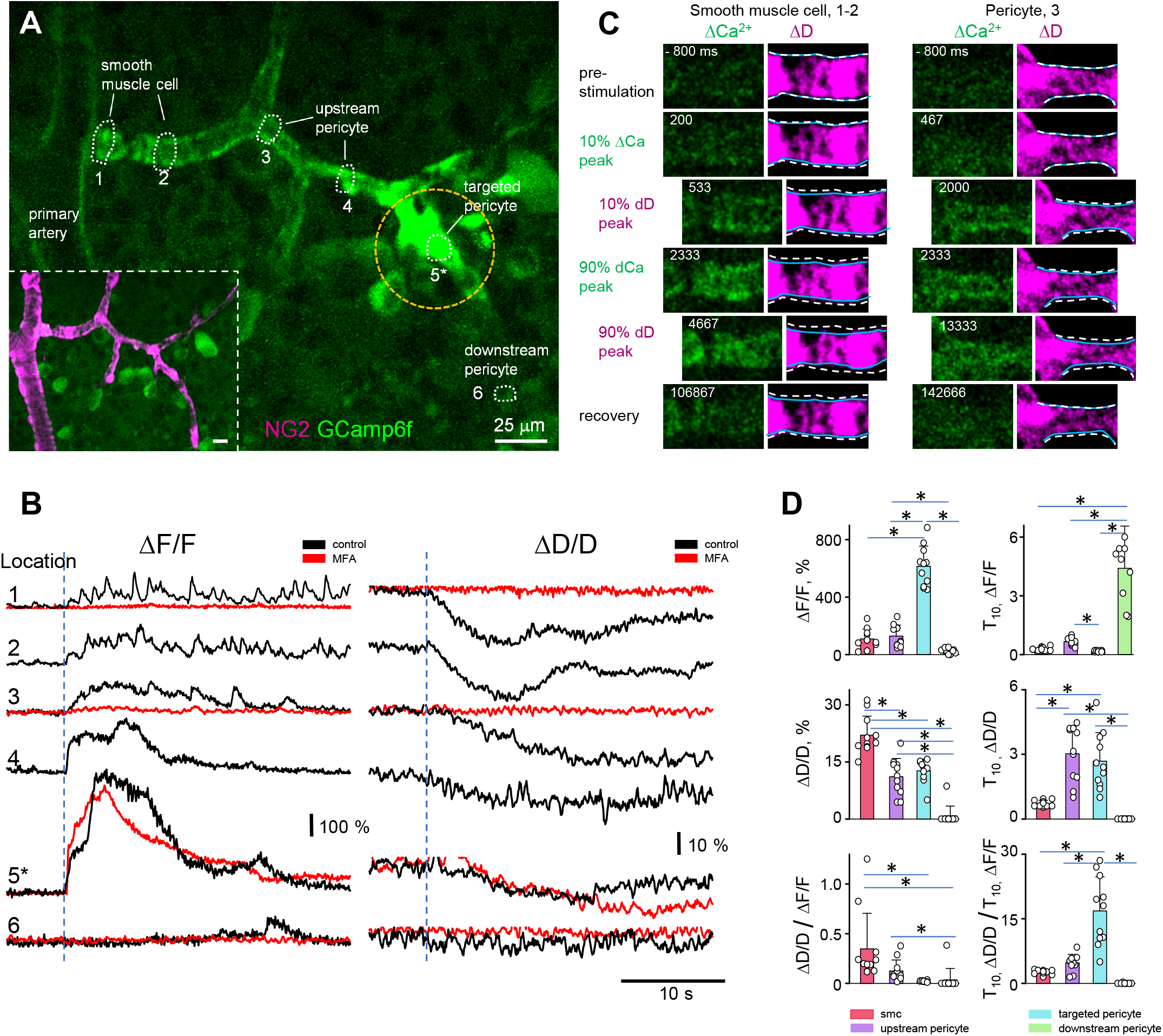
Vasomotor response is shaped by contractile cell-type specific temporal dynamics between Ca increase and vasomotor response. **(A)** Directional calcium increase in a vascular branch after local electric stimulation of a capillary pericyte (5) in an NG2-DsRed mouse. Dashed circle marks the border of local Muller cell activation. **(B)** Temporal kinetics of calcium raise and vasomotor response in contractile cells along the vascular branch shown in A. Calcium raise in non-targeted contractile cells and propagation of the vasomotor response was blocked by 40 μM meclofenamate (MFA, red); they were not affected in the targeted pericyte. **(C)** Remote smooth muscle cells at the artery have faster calcium raise and stronger vasomotor response than pericytes adjacent to the stimulation site. Pre-stimulation outlines of the blood vessels are shown by a dashed line in all frames while current outlines are marked by solid blue lines. **(D)** Calcium raise, vasomotor response, and Excitation-Constriction Coupling is significantly different between pericytes and smooth muscle cells. Data are shown as average ± SD; 11 samples, 6 mice, one-way ANOVA. *p<0.05

Finally, for each contractile cell type we determined the temporal relationship between Ca^2+^ increase and vasomotor response (Figure 5D). Ca^2+^ increase (ΔF/F) in the targeted pericyte was significantly higher than that in the other contractile cells (615 ± 139% in targeted pericyte; 129 ± 75% in upstream pericyte; 109 ± 68% in SMC; 27 ± 17% in downstream pericyte; 11 branches, 6 mice). Ca2+ increase in upstream pericytes was also significantly higher than in downstream pericyte. In the smooth muscle cells (SMCs), Ca^2+^ increase tended to be lower than in pericytes. However, the peak vasomotor response (ΔD/D), initiated by the Ca^2+^ wave was significantly larger in SMCs (22 ± 5% in SMC; 11 ± 4% in upstream pericyte; 12 ± 3% in targeted pericyte; 1 ± 1% in downstream pericyte; 11 branches, 6 mice). There was little or no vasoconstriction in the downstream pericyte. The efficiency of constriction, defined as the ratio of constriction to Ca^2+^ increase (ΔD/D / ΔF/F), was significantly higher in smooth muscle cell (0.35 ± 0.3 in SMC; 0.13 ± 0.11 in upstream pericyte; 0.02 ± 0.01 in targeted pericyte; 0.03 ± 0.1 in downstream pericyte; 11 branches, 6 mice). Temporal kinetics were also different between pericytes and smooth muscle cells. Time of 10% Ca^2+^ increase (T10ΔF/F) in the downstream pericyte, if detected, was significantly delayed (285 ± 90 ms in SMC; 661 ± 194 ms in upstream pericyte; 164 ± 35 ms in targeted pericyte; 4400 ± 2162 ms in downstream pericyte; 11 branches, 6 mice). Surprisingly when the time of vasoconstriction was compared (T10ΔD/D), it was significantly shorter in the remote smooth muscle cells than in the upstream pericytes adjacent to the targeted pericyte (706 ± 150 ms in SMC; 3030 ± 1283 ms in upstream pericyte; 2958 ± 1399 ms in targeted pericyte; 11 branches, 6 mice). When we adjusted time of vasoconstriction to the time of Ca^2+^ increase (T10ΔD/D / T10ΔF/F), smooth muscle cells were still significantly faster (2.6 ± 0.5 in SMC; 4.8 ± 1.9 in upstream pericyte; 19 ± 9 in targeted pericyte; 11 branches, 6 mice). Thus, higher ΔD/D / ΔF/F coupling efficiency in SMCs compared to pericytes may explain the earlier observations of faster response kinetics of vasomotor response in arteriolar branches during capillary stimulation.

### Diabetic retinopathy disrupts vascular connectivity map, impairing directionality and extent of vasomotor response

In this study we have demonstrated that the vascular cell coupling, propagation of Ca^2+^ increase and resultant vasomotor response rely on active GJs. We also show that these interactions are strengthened by and dynamically shift with changing sensory modality. In diabetic retinopathy (DR), GJs between endothelial cells and possibly endothelial to pericyte cells are selectively eliminated^6, 27^. We injected NG2-GCamp6f mice with streptozotocin (STZ) to induce Type 1 diabetes and to determine how selective elimination of GJs in the vascular relay affects cellular interactions and vasomotor response during the progression of DR. To validate an STZ-induced animal model, we measured the change in functional hyperemia, an early pathology in patients with diabetic retinopathy^28, 29^. We compared blood flow in the retinal capillaries *in vivo* in response to light flicker, as described earlier (Figure 6A and Suppl.Vid.2, 3)^30^. We found that blood flow was significantly reduced in STZ-treated animals compared to placebo-treated animals (Figure 6B,C; non-diabetic 44.8 ± 5.7 cells/s vs. diabetic 34.2 ± 8.4 cells/s; T-TEST, P < 0.001; 6 capillaries per animal, 5 mice per group). In freshly dissected retinal explants, we also found significant reduction in both pericyte-pericyte and pericyte-endothelial cell coupling compared to both sham controls and pre-diabetic animals (at the onset of hyperglycemia, ~3 weeks post STZ-injection (Figure 6D,E). This data is consistent with our earlier report, showing selective loss of Cx43-positive “strings” and diminished vasomotor response propagation^6^. A pharmacological block of GJs blocked all coupling of the targeted pericyte (Figure 6F), suggesting the progressive nature of DR-related impairment and partial preservation of GJ-mediated communication. Indeed bath application of SNP, partially restored DR-induced vascular coupling (Figure 6G). In contrast, application of SNP did not reverse the DR-induced loss of coupling directionality (Figure 7B).

**Figure 6.**
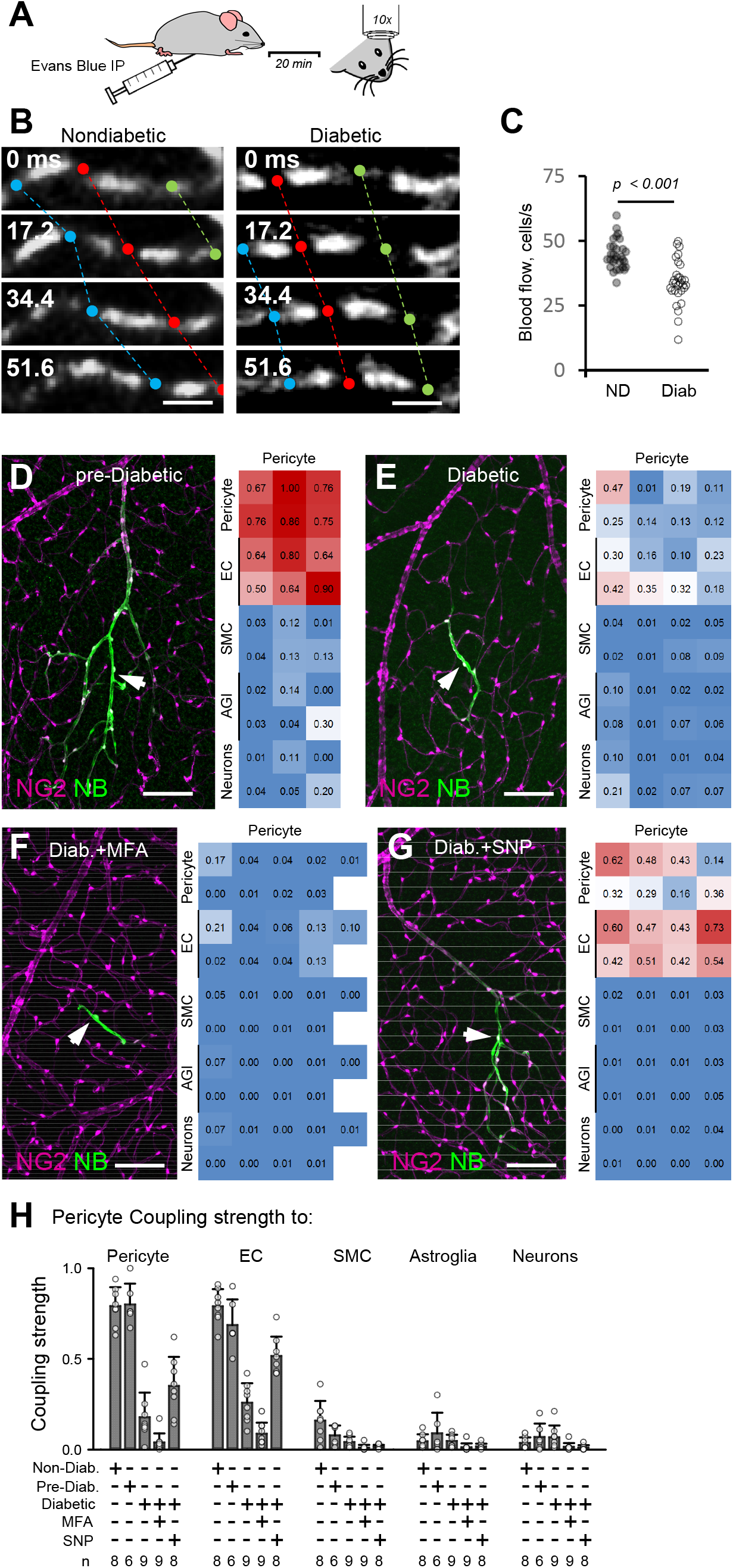
Functional connectivity maps are disrupted in diabetes. **(A)** Experimental paradigm of measuring retinal capillary blood flow in mice *in vivo*. **(B, C)** Assessment of retinal capillary blood flow *in vivo* reveals reduced blood flow in response to a light flicker (10 Hz) in mouse model of T1D. Evaluation of cell coupling in living retinal wholemount shows altered at the onset **(D)** and disrupted connectivity during T1D **(E)**. Coupling is further blocked in the presence of MFA (40 μM) **(F)**, but only partially rescued by NO donor (**G**, SNP 100 μM). **h.** Summary of the experimental data under each condition (n = animals per group).

**Figure 7.**
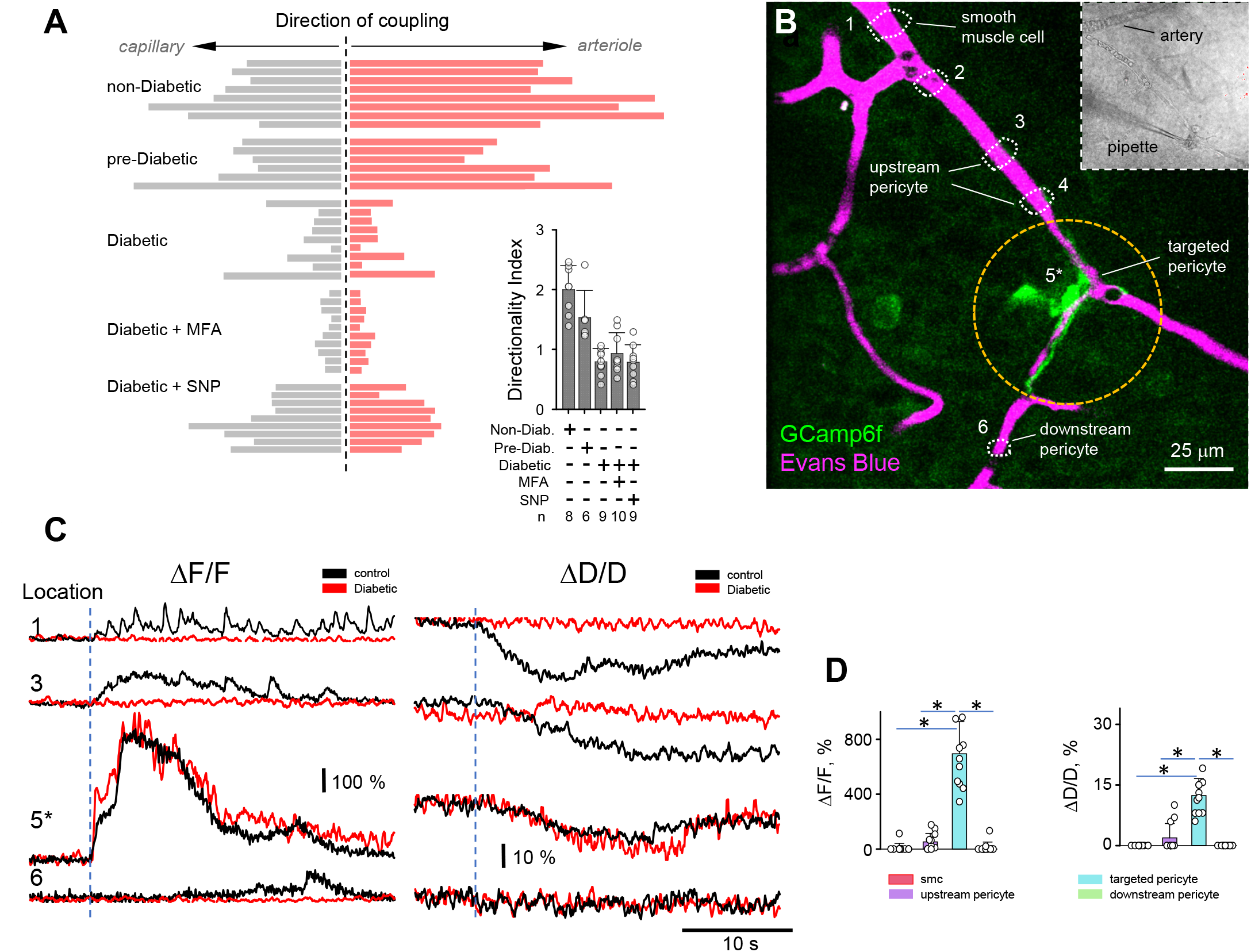
Vascular coupling, Ca signaling and vasomotor response are disrupted in diabetes. **(A)** In diabetic retina vascular cell coupling and its directionality was abolished. Global NO application enhanced cellular coupling but did not restore its directionality. **(B)** In living retinal wholemount of NG2-GCamp6f mouse, local depolarization of a pericyte by an electrode lead to local Ca raise and vasoconstriction. **(C)** Temporal kinetics of Ca raise and vasomotor response in contractile cells along the vascular branch shown in **A. (D)** Local Ca and vasomotor responses did not propagate along the vascular branch. Data are shown as mean ± SD; 12 samples, 6 mice, oneway ANOVA. *p<0.05

Next, in the living retinal wholemount, we focally stimulated a pericyte with a patch pipette while monitoring Ca^2+^ dynamics and vasomotor response (Figure 7B). To visualize blood vessel diameter, the mouse was intraperitoneally injected with Evans Blue to label plasma albumin and, therefore, blood (Figure 7B, magenta)^30^. Contractile cells were identified in transmitted light based on their characteristic “bump on the log” shape (Figure 7B). Stimulation of a targeted pericyte (location 5) induced Ca^2+^ increase, which did not propagate beyond the stimulated cell (Figure 7B,C). In line with the restricted Ca^2+^ wave, vasoconstriction was limited to the blood vessel underneath the stimulated pericyte (Figure 7B,C). Both Ca2+ wave and vasoconstriction at the targeted pericyte was similar between wild type and diabetic retinas (Figure 7D, wt: targeted pericyte ΔF/F = 615 ± 139%, ΔD/D = 12 ± 3%, n=11, 6 mice; diabetic: targeted pericyte ΔF/F = 693 ± 234%, ΔD/D = 12 ± 4%, n = 12, 6 mice; ΔF/F P = 0.35, ΔD/D P = 0.84, T-TEST). In contrast to wt, propagation of the calcium wave and vasoconstriction was significantly different in diabetic retinas (wt: upstream pericyte ΔF/F = 129 ± 75%, ΔD/D = 11 ± 5%, n = 11, 6 mice; diabetic: upstream pericyte: ΔF/F = 52 ± 62%, ΔD/D = 2 ± 4%, n = 12, 6 mice; ΔF/F P = 0.013, ΔD/D P < 0.001, T-TEST; wt: SMCs ΔF/F = 109 ± 68%, ΔD/D = 22 ± 5%, n = 11, 6 mice; diabetic: SMCs ΔF/F = 9 ± 33%, ΔD/D = 0 ± 0%, n=12, 6 mice; ΔF/F P < 0.001, ΔD/D P < 0.001, T-TEST). The absence of propagation in the downstream direction was similar in wt and diabetic retinas (wt: downstream pericyte ΔF/F = 29 ± 16%, ΔD/D = 1 ± 3%, n = 11, 6 mice; diabetic: downstream pericyte: ΔF/F = 13 ± 38%, ΔD/D = 0 ± 0%, n = 12, 6 mice; ΔF/F P = 0.22, ΔD/D P = 0.31, T-TEST). Thus, diabetic retinopathy caused disruption of communication among the vascular cells similar to the effect of GJ-blocker in wt.

## Discussion

In this work we describe a functional connectivity map of pericytes and endothelial cells that mediate the spatial and temporal precision of neurovascular signaling in the retina. We found that pericytes formed a precise 3-D mosaic across the retina. However, in response to sensory stimuli, the vasomotor activity propagated asymmetrically along the feeding branch. This directionality was observed when the stimulus matched neuronal receptive field center size and was driven along a highly discriminatory GJ-mediated relay between pericytes and endothelial cells. Pericytes and endothelial cells connected predominantly to other upstream neighboring pericytes and endothelial cells, and less to arteriolar smooth muscles, and not to surrounding neurons and glia. Below, we discuss the implications of our findings, potential mechanisms and new questions that may arise.

### Connectome of the neurovascular unit

Functional hyperemia is thought to rely on coordinated activity across a broad vascular network - sensing activity changes and then directing blood to an active region. With the exception of a few recent reports^8, 31^, the mechanistic studies of spatial interactions during functional hyperemia have mainly focused on correlations between neural activity and vasomotor elements within a restricted region.

Here, in the living retina, we mapped pericytes across a broad capillary network to reveal that they form a mosaic. Interestingly, the observed mosaic was not simply among the pericytes on the same vascular branch, but also among pericytes on disparate capillaries. Our analysis indicates that this arrangement was not a result of inherent isotropy of the vascular network itself, but rather is an evidence for the optimized activity sensing across the tissue volume. However, such a “symmetric” structure, poses a challenge: how to discriminate an active region from its quiescent neighbor? We showed that the spatial accuracy of the vasomotor response was mediated by discriminatory connections among vascular cells along the feeding vascular branch. We found that pericytes and endothelial cells form a functional relay unit by coupling to each other, and not to surrounding neurons and glia. This is intriguing given the abundant expression of various connexins across a wide range of cells in the retina^13^. While the regulatory mechanisms for this discriminatory connexin expression remain unclear, this cellular specificity may provide an important insight into the nature of the vascular signaling and its accuracy. Furthermore, limited coupling between smooth muscle cells of the arterioles may have implications to containing the spread of vasoactive signal past the activity site, a step needed to avoid a non-specific broad blood supply. The spatially contained spread of vasoactive signal during functional hyperemia supports the presence of local functional domains^8^.

### Spatial tuning of neurovascular coupling

Our findings also suggest that vascular signaling is not only capable of spatially precise delivery of blood to the active region, but is optimized to focal stimulation. The strengthening of both vascular cell coupling and vasomotor response during focal stimulation is likely driven by an optimized neuronal response during the neuronal receptive field center stimulation. This finding is significant as it establishes an upper resolution of fMRI BOLD imaging at the level matching the receptive field of neuronal interactions. It also provides a mechanistic basis for *in vivo* observations of a spatial resolution at 100–400 μm, corresponding to the areas of cortex activated by a whisker stimulation or by a single ocular dominance column in the visual cortex^32–35^. Such spatial precision of vasomotor response would require recruitment of the contractile cells exclusively along the active vascular branch and only up to the active site. We found that the recruitment of contractile cells along the active branch was mediated through the discriminative coupling of the vascular cells. This GJ-mediated coupling was essential for fast propagation of a Ca^2+^-wave leading to a vasomotor response along that vascular branch. Ca^2+^-wave was restricted to the vascular cells without involvement of neurons, astrocytes, or Müller cells, consistent with a recently described vascular relay circuit^13^. It was likely spreading through ECs, as the current spread through the microvascular endothelium has much higher efficiency than between pericyte to pericyte and pericyte to EC^14, 17, 36, 37^. This fast and non-decaying signal propagation through ECs may rely on active regenerative mechanism^38^ and allowed nearly instantaneous activation of cells along the active vascular branch. Interestingly, we showed that both Ca^2+^ dynamics and excitation-contraction coupling was most efficient in SMA, which may explain faster vasomotor response on remote arterioles compared to capillaries shown both *in vitro^36^* and *in vivo*^8^.

Once appropriate vascular branch has been activated, the next task is to restrict signal propagation to the active branch without affecting other areas that are supplied by the same feeding artery. First, our cellular tracing and GCaMP6f experiments using intact retina revealed that vascular cells were electrotonically insulated from the surrounding glia and neurons, thus limiting radial spread. Second, pericytes and ECs coupled strongly along the capillary and weakly to ECs and smooth muscle cells on the arteriole. This is consistent with the earlier electrotonic studies in isolated vascular branches where the conductance dropped abruptly at the branching point^14^. Extending recent reports, we also showed that the vasomotor signal was observed from the upstream arteriole up to the active side^8, 16, 39^. Consistently, vascular coupling, Ca^2+^-wave and vasomotor responses were recorded on the upstream side of the active region. This directional response towards the feeding artery was driven exclusively by local NO production and was lost during global sensory stimulation or bath NO application. We further hypothesized that this NO-mediated directionality depended on preferential opening of ECs’ GJs in the upstream direction. ECs are highly polarized with respect to blood flow^40, 41 42^. Specifically, caveolae with caveolin-1 and eNOS are abundant in the upstream end of ECs^43^. Caveolin-1 inhibits the activity of eNOS. Upon shear stress^44^ or Ca^2+^ elevation (as in our zap experiments), eNOS leaves caveolin, binds to Ca^2+^-calmodulin and produces NO^45^. Thus, NO production appears to be compartmentalized^46^ in the upstream end of ECs. Consistent with our experiments, NO was shown to nitrosalate Cx43 GJs resulting in their opening^47, 48^. While further studies are needed, polarized distribution of caveolin-1-eNOS complex in EC may be a mechanism to control directionality and spatial precision of vasomotor response via selective activation of GJs.

### Implication to neurovascular pathology in diabetic retinopathy

In diabetes, vascular pathology is associated with high glucose, causing activation of protein kinase C pathway^49–51^ and reduced activity of Cx43^6, 12, 27, 52, 53^. In our experiments with the retina of diabetic mice, cellular coupling, vasomotor response and blood flow were impaired. This is consistent with reduced flicker response and abnormal blood flow in the retina of diabetic patients^53–55^. As we showed in a mouse model of diabetic retinopathy, expression of the Cx43 GJ along the vascular relay is preferentially downregulated^6, 13^. Downregulation of GJs lead to restriction of Ca2+ wave and vasomotor response experiments along the vascular branch consistent with a ~5-fold increase in voltage decays along the retinal microvasculature in rat model^56^. Disrupted vascular cell connectivity and reduced responses to vasoactive signals may contribute to decline in functional hyperemia observed early in the disease.

## Acknowledgments

This work was supported by NIH grants R01-EY026576 and R01-EY029796 (BTS). The authors thank Dr Rajiv Ratan, MD, PhD for comments on the manuscript.

## Author Contributions

TK-O, E.I., P.B. and B.T.S. designed, conducted the experiments and wrote the paper.

## Material and Methods

In all experimental procedures, animals were treated in compliance with protocols approved by the Institutional Animal Care and Use Committee (IACUC) of Weill Cornell Medicine (WCM), and in accordance with the National Institutes of Health Guide for the Care and Use of Laboratory Animals. The use and application of streptozotocin was in accordance with safety protocols approved by WCM’s Environmental Health and Safety (EHS), Institutional Biosafety Committee (IBC) and IACUC Protection and Control sub-committee (P&C).

### Experimental animals

Diabetes was induced in three mouse lines: C57BL/6 mice (Jackson Laboratory, Stock #: 000664, RRID:IMSR_JAX:000664), NG2-DsRed mice (Jackson Laboratory, Tg(Cspg4-DsRed.T1)1Akik/J, Stock #: 008241, RRID:IMSR_JAX:008241) and NG2-Cre-GCamp6f, generated by crossing FVB-lfi208Tg(Cspg4-cre)1Akik/J (Jackson Laboratory, Stock #: 008533, RRID:IMSR_JAX 008533) with Ai95(RCL-GCaMP6f)-D loxP (Jackson Laboratory, Stock #: 024195, RRID:IMSR_JAX 024195). We used the streptozotocin (STZ) diabetic mouse model^57^. Male mice aged 6 to 8 weeks were fasted for 4 hours prior to the injections. The animals were injected intraperitoneally on five consecutive days with 50 mg/kg STZ (Sigma-Aldrich, S0130) freshly dissolved in citrate buffer (pH 4.5). Control animals received a citrate buffer injection without STZ. In our STZ mouse model of diabetes, the levels of blood glucose reached maximum elevation at 1 month after STZ injection and remained elevated. The diabetes was defined by non-fasting blood glucose greater than 300 mg/dL verified 1 month after the last STZ injection and confirmed on the day of the experiment.

### Retinal wholemount preparation

Methods for wholemount tissue preparation have been described in detail previously^58^. After the animal was euthanized, its eyes were enucleated and placed in bicarbonate-buffered Ames solution (Ames; Sigma, A1420), equilibrated to pH7.4. It has been shown that variations in O2 level in brain tissue can affect functional hyperemia^5^. To reduce any discrepancy O2 level was maintained between 19-24%, checked with oximeter (WPI ISO2-D) in the chamber. After dissection of the eyes cornea, iris, and lens were removed. The retina was dissected into four equal quadrants and attached photoreceptor surface down on a modified Biopore Millicell filter (Millipore). This preparation was transferred to a recording chamber and bathed (1 ml/min) with Ames. Pharmacological agents were also prepared in Ames. All experiments were performed at a near physiological temperature of 32°C.

### Identification of pericytes and mosaic measurements

In the retina wholemount, identified pericytes were targeted on capillaries in the superficial vascular layer, avoiding those located on arterioles and veins. Capillaries were defined based on several morphological criteria: (i) diameter not exceeding 10 μm, approximately equivalent to the diameter of red blood cells, which are readily present in the living tissue, (ii) lack of a smooth muscle actin. Initially, we targeted fluorescently labeled pericytes in NG2-DsRed mice. In genetically unmodified mice, we identified pericytes based on “bump on a log” appearance of the individual pericytes on the abluminal side of the vessel wall (Kawamura, Sugiyama et al. 2003), or by using 10-minute incubation in NeuroTrace 500/525 Green Fluorescent Nissl Stain (1:200 in HEPES-Ringer)^20, 30^. We measured the nearest neighbor distances (NND) and shortest route along vessel (SRAV) between pericytes using ImageJ (NIH, USA). NND was defined in an arrangement of collapsed optical slices within each vascular lamina. We measured N=275 NNDs between pericyte somas on the collaps ImageJ, Simple neurite tracer in a 3D reconstruction. We made N=141 measurements from all three layers. The SRAV routes were manually checked for errors, less than 2% had to be corrected (e.g. manually link non-touching pericyte-endfeet). Normal distribution of datasets for mosaic arrangement had been tested with Excel descriptive statistics kurtosis (<1) and skewness (<1).

### Cellular coupling and directionality assessment

Cell coupling was assessed using Neurobiotin probe, as described^13^. Electrodes were pulled from borosilicate glass (1B150F-4) with a P-97 Flaming/Brown puller and had a resistance of ~1-2 MΩ. First, the pipette was filled with filtered Ames solution supplemented with 100 μg/ml papain (~1 Unit/ml). To dissolve the vascular basement membrane covering pericytes, the papain-containing solution was focally applied around the target cell for 5 min. Second, a fresh pipette was filled with intracellular solution containing (in mM): 120 Cs-gluconate, 10 tetraethylammonium chloride (TEA-Cl), 1.0 CaCl2, 1.0 MgCl2, 11 ethylene glycol-bis(beta-aminoethyl ether)-N,N,N’,N’-tetraacetic acid (EGTA), and 10 sodium N-2-hydroxyethylpiperazine-N’-2-ethanesulfonic acid (Na-HEPES), adjusted to pH 7.2 with CsOH. The solution was supplemented with 2% Neurobiotin (Vector, SP-1120) and 0.5% Alexa488-hydrazide (ThermoFischer Scientific, A10436). This new pipette was pressed against the cleaned target and the cell membrane was gently pulled inside the pipette until a >200 MOhm seal was established. The alternating voltage steps between −300 mV and +50 mV, 2 Hz, were applied for 1 min to confirm successful targeting of a cell after filling with Alexa488-hydrazide. If the target cell was selectively backfilled with Alexa488, Neurobiotin was electroporated for additional 3 min using +200 mV −50 mV, 2 Hz voltage steps. If either the targeted cell was not filled with Alexa488 or Alexa was detected outside of the injected cell, the preparation was discarded. Following Neurobiotin electroporation, the preparation was left for 15 min to allow intercellular Neurobiotin diffusion. All electroporations were made with MultiClamp 700B patch-clamp amplifier (Molecular Devices, Sunnyvale, CA) using Signal software (CED, UK). We have limited injecting cells within the superficial layer for the following reasons: a) a robust access to all cell types for both tracing and targeted stimulation, without causing physical damage to the wholemount tissue and thus substantially reducing the possibility for non-specific tracer pickup, b) all blood vessel types, from the arteriole to finest capillaries, are present in the superficial layer, allowing for the coupling assessment along a contiguous vascular tree, and c) diverse cell types are present, including astroglial cells, that are known to play an active role in neurovascular coupling^7^. The Neurobiotin coupling was assessed across all vascular layers (Extended Data S2).

For each targeted cell type, a coupling strength was measured as a ratio of the Neurobiotin stain fluorescence intensity in cell bodies of six nearest neighbors to the Neurobiotin stain intensity in the injected cell body. Therefore, the values approaching 1 indicate strong coupling, while 0, such as in the presence of GJ-blocker MFA, no coupling (Figure 2F). In case of pericytes and endothelial cells, Neurobiotin stain intensity was measured in three cells in each direction up- and downstream the vascular branch from the injected cell (Figure 2E). When heterocellular coupling was detected, the coupling strength was measured for each cell type combination to yield a coupling matrix (Figure 2G). The directionality of both vascular cell coupling and vasomotor response propagation was assessed using a directionality index (DI), a ratio between magnitude of upstream vs. downstream bias. DI values >1 indicate a bias towards the feeding artery. This simple approach provided a robust and reproducible assessment of the response directionality bias and was not intended to discriminate between pericytes and endothelial cell involvement.

### Vasomotor response induction and quantification

Pericytes were focally stimulated under upright Nikon FN1 microscope by a current pulse (7 μA, 2 ms; Grass Technologies) using an electrode filled with Ames solution. Electrodes were pulled from borosilicate glass (WPI, 1B150F-4) with a P-97 Flaming/Brown puller (Sutter Instruments, Novato CA) and had a measured resistance of 3-5 MΩ. For consistency across all experiments, the electrode was placed near the cell body of the targeted pericyte. During focal “puff” stimulation, electrode solution was supplemented with a vasoactive compound and delivered with picospritzer (Parker Hannifin) via broken patch pipette positioned above the targeted pericyte. For the light stimulation experiments, the microscope’s illuminator was used to deliver spot of light that was centered on the targeted pericyte cell body and focused on the photoreceptor cell layer. The tissue was adapted at 30 cd/m2, and stimulus was 270 cd/m2. Light spot flicker (40 μm diameter, 10 Hz) was controlled by a shutter (Uniblitz, Vincent Associates). Responses to stimuli were captured on video or time-lapse photo with a microscope-mounted Sony A7s full frame camera. Images of blood vessels were analyzed in ImageJ, using a region of interest (ROI) tracing tool. At each experimental condition, the capillary lumen cross sections were mapped at 2 μm steps along the capillary.

### Two-photon assessment of calcium dynamics, vasomotor response and blood flow

Retinal quadrants attached to a filter were transferred to a recording chamber on the stage of an upright ThorLabs Bergamo II two-photon microscope with a tunable femtosecond TI sapphire laser. GCamp6f, Evans Blue and DsRed were simultaneously excited at 920 nm. Evans Blue was used to visualize blood vessels and was i.p. injected in the living mouse 30 min prior to dissection^30^. Vasomotor response was induced by an electrode as described above. Movies were taken at 15 fps rate. All data were analyzed in ImageJ.

For the blood flow assessment, intraperitoneal injection of 100 μl (100 μg/ml) Evans Blue was done 30 min prior to measurements to visualize blood plasma. The animal was anesthetized with a mixture of 150 mg/kg ketamine and 15 mg/kg xylazine. The pupils were dilated with 0.5% tropicamide ophthalmic solution, and a coverslip was placed on each eye with GONAK ophthalmic solution. Mice were mounted with SG-4N mouse head holder (Narishige) on an upright ThorLabs Bergamo II two-photon microscope. Blood flow was measured under 10X super apochromatic objective with 7.77 mm working distance and 0.5 NA (TL10X-2P, ThorLabs, Newton, NJ). Evans Blue was illuminated with 920 nm wavelength and the measurements were taken at 116 – 400 frames per second rate. Blood flow was estimated as number of blood cells passing through a capillary per second.

### Immunohistochemistry

After vasomotor assessment, or pericyte tracing each sample still attached to the Biopore insert was submersion-fixed in freshly prepared fixative (0.25% PFA, 4% carbodiimide in Phosphate-buffered saline - PBS) for 15 minutes at room temperature. The fixed samples were washed in PBS and the retinas were separated from the insert. We visualized NB overnight with streptavidin-A488, mixed with To-Pro-3 Iodine (1:30000; far red, T3605, Thermo Fisher Scientific). In multi-labeling experiments, wholemounts were incubated in a mixture of primary antibodies, followed by a mixture of secondary antibodies. Retinal wholemounts were blocked for 10 h in PBS, containing 5% Chemiblocker (Chemicon), 0.5% Triton X-100, and 0.05% sodium azide (Sigma, St. Louis, MO). Primary antibodies were diluted in the same solution and applied for 72 h, followed by incubation for 48 h in the appropriate secondary antibody, conjugated to Alexa 488 (1:1000; green fluorescence, Molecular Probes), Alexa 568 (1:1000; red fluorescence, Molecular Probes), or Cy5 (1:500; far-red fluorescence, Jackson). All steps were carried out at room temperature. After staining, the retinal pieces were flat mounted on a slide, ganglion cell layer up, and coverslipped using Vectashield mounting medium (H-1000, Vector Laboratories). The coverslip was sealed in place with nail polish. To avoid extensive squeezing and damage to the retina, small pieces of a broken glass coverslip (number 1 size) were placed between the slide and the coverslip. The primary antibodies used in this study were the following: rabbit anti-Cx43 (Cx43, 1:2000, Sigma-Aldrich, C6219, RRID:AB_476857), rabbit anti-NG2 coupled to Cy3 fluorescent label (NG2, 1:500, EMD Millipore, AB5320C3, RRID:AB_11214368). Retinal samples were imaged under a Nikon Eclipse Ti-U confocal microscope (Morell Inst., Melville, NY). The samples were imaged under identical acquisition conditions, including: laser intensity, photomultiplier amplification, and Z-stack step size. All images were processed and analyzed using ImageJ. In total, we evaluated 1007 individual pericytes in non-diabetic and 561 pericytes in diabetic animals.

### Statistical Analysis

Statistical analysis was performed in SigmaPlot 14 (Systat, RRID:SCR_003210). For multiple comparisons, analysis of variance (ANOVA) with Tukey’s post-hoc, or repeated measures ANOVA were used. The data are presented as mean ± SD. N is the number of animals per group. To avoid introduction of non-independent data into statistical analysis, first, multiple samples were averaged within animal, then the data between animals were compared^59^.

---

**Suppl. Figure 1.**
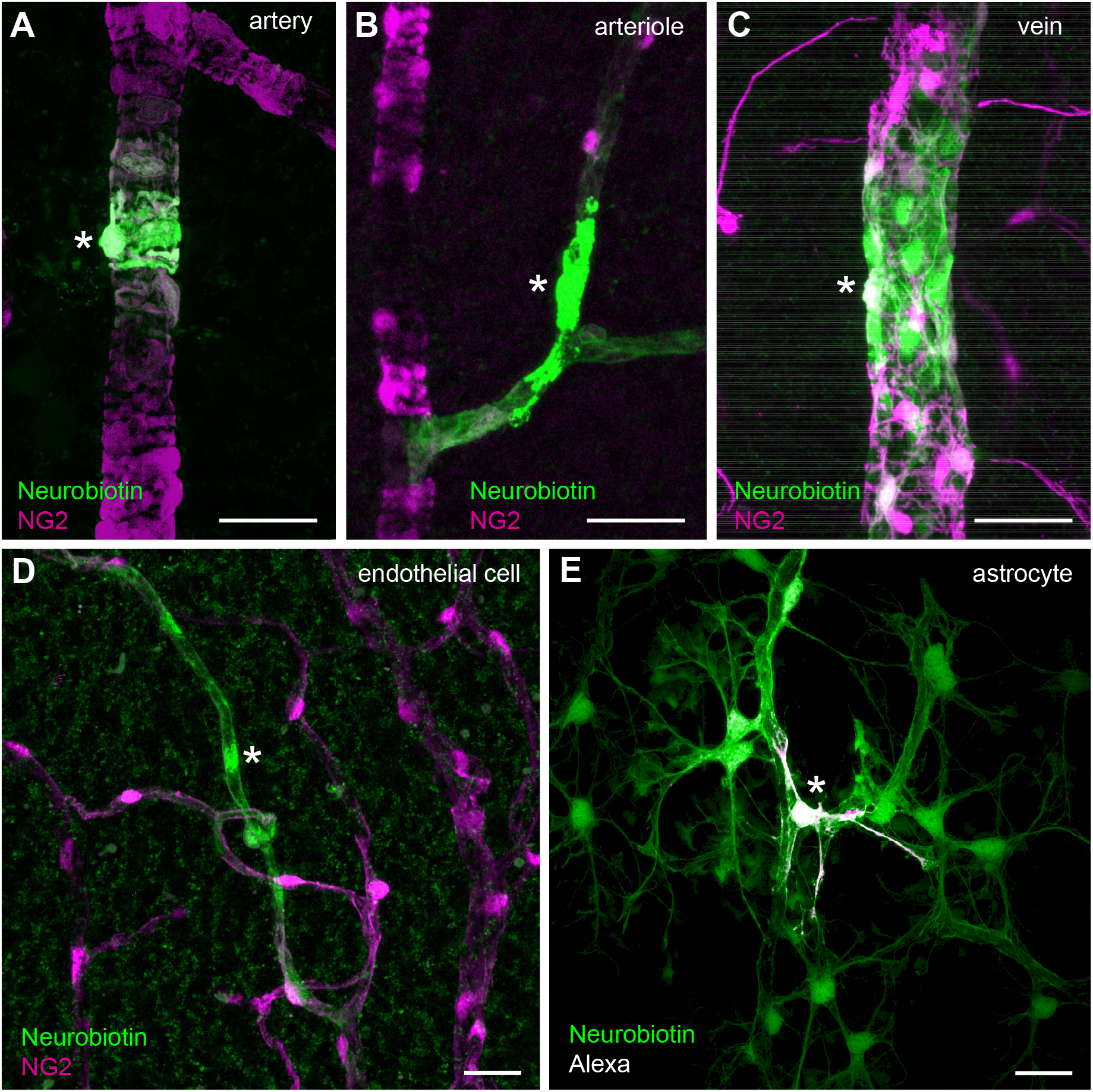
Gap-junction mediated coupling patterns across pericytes, smooth muscle cells, astrocyte and endothelial cells. **(A)** Representative tracing results following arterial and **(B)** arteriolar terminal smooth muscle cell injection with Neurobiotin probe (green) in the NG2-DsRed mouse retina. **(C)** Mural cell coupling pattern in the retinal vein **(D)** Representative tracing results following targeted Neurobiotin injection into endothelial and **(E)** astroglial cells. *injected cell. Scale bar: 50 μm.

